# Characterization of the adult zebrafish electrocardiogram

**DOI:** 10.1101/2022.02.02.478776

**Authors:** E. Arel, L. Rolland, J. Thireau, AG. Torrente, E. Bechard, J. Bride, C. Jopling, M. Demion, J-Y. Le Guennec

**Affiliations:** PhyMedExp, Université de Montpellier, Inserm U1046, UMR CNRS 9412, Montpellier, France; IGF, Université de Montpellier, Inserm U1191, UMR CNRS 5203, Montpellier, France

**Keywords:** Electrocardiogram, Hyperosmotic therapy, Hypothermic therapy, longitudinal studies, Bazett’s formula, zebrafish

## Abstract

**Background:** The use of zebrafish to explore cardiac physiology has been widely adopted within the scientific community. Whether this animal model can be used to determine drug cardiac toxicity *via* electrocardiogram (ECG) analysis, is still an ongoing question. Several reports indicate that the recording configuration severely affects the ECG waveforms and its derived-parameters, emphasizing the need for improved characterization.

**Methods:** ECGs were recorded from adult zebrafish hearts in 3 different configurations (unexposed heart, exposed heart and extracted heart) to identify the most reliable method to explore ECG recordings at baseline and in response to commonly used clinical therapies.

**Results:** We found that the exposed heart configuration provided the most reliable and reproducible ECG recordings of waveforms and intervals. We were unable to determine T-wave morphology in unexposed hearts. In extracted hearts, ECG intervals were lengthened and P-waves were unstable. However, in the exposed heart configuration, we were able to reliably record ECGs and subsequently establish the QT-RR relationship (Holzgrefe correction) in response to changes in heart rate.

**Conclusions:** The exposed heart configuration appears to be the most reliable technique to record ECGs in adult zebrafish. In this configuration, the QT-RR relationship, an important parameter in cardiac toxicity evaluation, can be determined using the Holzgrefe correction.

## Introduction

The electrocardiogram (ECG) is a widely used technique for analyzing the electrical activity of the heart and provides a wealth of information enabling clinicians to determine discrepancies in patients suffering from a variety of different cardiomyopathies. ECG is also a standard control used during drug development and current legislation stipulates strict ECG criteria which must be met before a novel drug can be approved. The typical ECG consists of 3 main components, the P wave, which represents the depolarization of the atria, the QRS complex, which represents the depolarization of the ventricles occurring during systole and the T wave, which represents the ventricular repolarization occurring during diastole (Figure 1A–B). One commonly assessed ECG parameter is the QT interval (the time interval between the start of QRS complex (ventricular depolarization) and the end of the T wave (ventricular repolarization). Prolongation of the QT interval may result in fatal arrhythmias which is why this parameter is a standard measurement when testing novel drug cardiotoxicity. New drugs must not significantly affect the QT interval in order for them to be approved for use in humans. Because the QT interval is inherently linked to the heart rate it is standard practice to correct the QT interval to take into account changes/differences in heart rate which are unrelated to the QT interval. Such changes would otherwise confound this measurement (for example someone with a slow heart rate will have a longer uncorrected QT interval than someone with a fast heart rate when they are in fact the same) resulting in the corrected QT or QTc. Prior to expensive clinical trials it is important to first assess cardiotoxicity in animal models such as zebrafish. Zebrafish cardiac physiology is highly comparable with humans and in this respect the zebrafish has become a powerful tool for determining the cardiotoxicity of novel pharmacological agents [1–6]. In particular, humans and zebrafish both exhibit comparable ECGs [1,2,7,8] with discernable P waves, QRS complexes and T waves.

**Figure 1:**
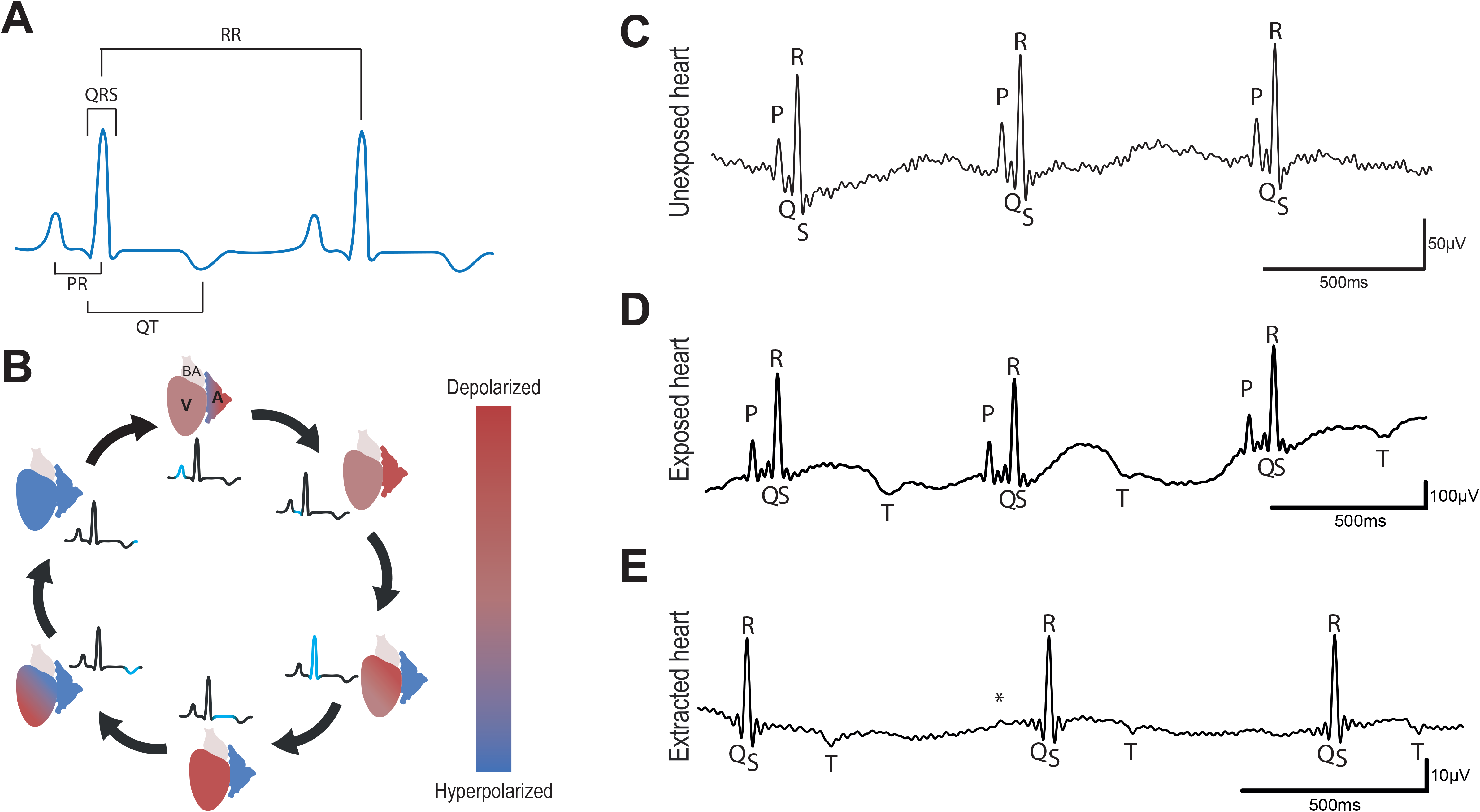
ECG representation and typical recordings **A**, Schematic representation of the interval measurement method. We chose to take the peak of the P and T waves to determine PR and QT intervals respectively to reduce the uncertainty of the measurements when taking before P waves or after T waves; **B**, Cartoon representing the depolarization and repolarization of the zebrafish heart. A: atrium, V: ventricle, BA: bulbus arteriosus; **C**, unexposed heart; **D**, exposed heart; **E**, extracted heart, * indicates putative P wave. Traces show three consecutive ECG complexes after application of a 50Hz low-pass filter.

The first ECG recordings in zebrafish were obtained by Milan *et al.* [1]. Subsequently, several research groups have tried to adapt the recording techniques and/or analysis methods to improve the quality and reliability of the signal. Despite some improvements, Liu *et al.* [4] highlighted discrepancies between the ECG signals obtained from different research groups regardless of their recording methods. Currently, there are two different techniques used by research teams to record ECG in adult zebrafish hearts. The first is non-invasive and involves positioning the electrodes on the body surface [9–11], while the second technique involves inserting the electrodes either 1mm into the dermis (*i.e.* in the pectoral muscles) or directly onto the surface of an exposed heart [1,3–5,12–18]. It is apparent that the choice of technique has a major influence on the raw ECG signal. For example, using the non-invasive technique it appears that Q, S and T waves are difficult to identify [9,11], and are generally assigned manually after the raw data has been processed [10]. In contrast, by using electrodes that allow direct access to the hearts electrical activity it is possible to reliably identify P waves, the Q, R, S complex and T waves [2–5,15,18,19]. Furthermore, post-experimental mathematical processing can also be used to reduce background noise [17,20–22]. However, it should be noted that T waves generated during the repolarization of the ventricle are difficult to discriminate using either technique. T wave analysis is an important factor in determining the QT interval during cardiotoxicity testing and in this sense it is important to be able to reliably and accurately measure this parameter. At present there appears to be confounding data regarding the nature of T waves in the adult zebrafish heart. For example, although several studies have reported negative T waves [1,10,14,23], other groups have actually recorded positive T waves [2–4,11–13,15,18,19]. It should be noted that in some cases, the T wave appears to be rather difficult to identify distinctly when the signal is recorded using a microelectrode array [16,17,22]. Lastly, ECGs can also be recorded on isolated hearts [24,25]. However, in this configuration the interpretation of the results is confounded by the absence of autonomic regulation by the nervous system [25–27]. This is particularly relevant as recent evidence suggests that the cardiotoxicity of certain drugs linked to ventricular repolarization involves autonomic dysregulation [28,29].

In this study, we aimed to evaluate and compare ECG profiles, measured longitudinally on the same animal, in 3 different configurations (unexposed heart, exposed heart and isolated heart) in order to determine which technique is the most accurate. Furthermore, we endeavored to characterize the QT-RR relationship in each configuration in order to determine which of these techniques is the most accurate and reliable for assessing the QTc interval. To this end, we also sought to determine what effect clinical therapeutic treatments such as hyperosmolality and hypothermia (targeted temperature management) had on ECG recordings in the 3 different configurations.

## Material and Methods

### Zebrafish strains and husbandry

Zebrafish were maintained under standardized conditions and experiments were conducted in accordance with local approval (APAFIS#2021021117336492 v5) and the European Communities council directive 2010/63/EU. All experiments were performed on 6-8 month old AB wildtype fish.

### ECG recording

Zebrafish were anesthetized in tricaine (160mg/L). Then, they were placed ventral side up in a slit sponge. Two 29-gauge stainless steel micro-electrodes (MLA1213, AD Instruments) were positioned along the ventral midline. In unexposed heart configuration, the positive electrode was positioned just above the heart and the negative electrode in front of the anal fin to record ECG. In the exposed heart configuration, we surgically opened the cardiac cavity and the electrodes were positioned close to, but not touching, the cardiac muscle. Lastly in the extracted heart configuration, the heart was extracted under a dissection microscope and the electrodes were positioned in the main axis of the ventricle. In this configuration, the heart activity was stable during the ECG recording. In all conditions and in accordance with the 3Rs, ECGs were recorded for exactly one minute only. After recording ECG in the unexposed heart configuration, individual fish were returned to their tanks and maintained separately for few days before recording the ECG again in the exposed heart configuration. Individual fish were again allowed to recover for a few days in the system before recording ECGs in the terminal extracted heart configuration. In the three configurations, ECG signals were amplified and digitized using a BioAmp (FE231, AD Instruments) and a PowerLab (16/35, AD Instruments). Data were subsequently processed using LabChart Pro v8 Software and the ECG analysis module (AD Instrument). Recordings were made in the range 0-10mV. A 50Hz notch digital filter was then applied and a sliding averaging algorithm provided by the software was used to smooth the traces. To reduce the biological variations, the same animals were used in the three different configurations of ECG measurements.

### QT analysis and correction

ECG were manually analyzed as auto-identification waveforms is often associated with cursor placement errors. The most common software error was a failure to distinguish between P and R waves resulting from excessive noise or the unstability of the isoelectric line. As a result, the software also failed to automatically and reliably identify the Q wave. Thus, all ECG waveforms were systematically identified and the errors in positioning the PQRST cursors were fixed by operator to avoid miss-interpretation. We chose to use the Ppeak, Qpeak, Speak and Tpeak to analyze the ECG instead of the beginning/end of the waveforms in order to reduce both the risk of miss-placement of the wave and the inter-operator dependence of the results.

The PR interval was estimated as the time between the peak P and the peak R waves and the QT interval as the time between the peak Q and peak T waves (Figure 1A).

Due to interspecies variability of the QT-RR relationship, we used the corrected QT (QTc) formula described by Holzgrefe *et al.* (2014) [30]:

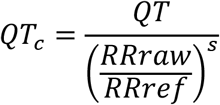

*s* is the slope of the linear relationship Log(QT)=f(Log(RR)) and RRref is the reference RR. The beating frequency is around 120 beats per minute thus RRref is equal to 0.5s [1,3].

### Solutions and experimental protocols

Isolated hearts were kept in a Tyrode solution: NaCl 140mM, KCl 5mM, CaCl_2_ 2mM, MgCl_2_ 1mM, HEPES 10mM, pH 7.4 (309mosm.L^−1^). To determine the effects of reducing the temperature, the Tyrode solution was chilled from 28°C to 5°C. One-minute ECG recordings were obtained in warm Tyrode (control, 28°C), cold Tyrode (5°C) and warm Tyrode as a washout condition.

To produce an osmotic shock, we first prepared an isotonic hypo-Na Tyrode solution: NaCl 90mM, mannose 100mM, KCl 5mM, CaCl_2_ 2mM, MgCl_2_ 1mM, HEPES 10mM, pH 7.4 (309mosm.L^−1^). We then made a hyposmotic solution: NaCl 90mM, KCl 5mM, CaCl_2_ 2mM, MgCl_2_ 1mM, HEPES 10mM, pH 7.4 (209mosm.L^−1^). Finally, we prepared a hyperosmotic solution: NaCl 90mM, mannose 200mM, KCl 5mM, CaCl_2_ 2mM, MgCl_2_ 1mM, HEPES 10mM, pH 7.4 (409mosm.L^−1^). The ECGs were recorded in all conditions following the sequence: Tyrode-Isotonic-Hyposmotic-Isotonic-Hyperosmotic-Isotonic.

### Statistical analysis

Data were expressed as mean (SEM). The parameters measured (PR, QRS and QT intervals, HR and QTc) were compared between the different recording configurations first and then within each configuration for the temperature and osmotic protocols. For each recording, PR and QT intervals were excluded when P or T waves were only detected in 15% or less of the ECG complexes. We used repeated measures 1-way ANOVA followed by Tukey’s *post hoc* test to compare 3 or more normal groups, or Kruskal-Wallis test followed by Dunn’s *post hoc* test to compare 3 or more groups which did not pass normality testing. To compare only 2 paired groups, we used students paired t tests or its non-parametric equivalent Wilcoxon’s test according to the normality of the values. p<0.05 was considered statistically significant.

## Results

### Basal characteristics of adult zebrafish ECG

In order to determine the most reliable method for analyzing the QT interval in adult zebrafish we recorded ECGs in 3 different configurations, unexposed heart, exposed heart and extracted heart (Figure 1C–E). From these recordings we were able to determine the average PR interval, QRS duration, RR interval, heart rate, QT interval and calculate the QTc interval (Figure 1A and Table 1). Our analysis indicates that there is no significant difference in heart rate between the three configurations (Table 1). When we compared the two *in vivo* configurations (unexposed heart and exposed heart) we found that the QRS interval was longer in the exposed heart configuration suggesting a possible slowing of ventricular conduction. However, assessing the QT interval in the unexposed heart configuration presented difficulties as we were only able to detect the T wave in 8 out of 20 samples (Tables 1 and 2, supplemental Tables 1 and 2). Under normal conditions in humans, T waves are primarily positive, however, although negative (inverted) T waves are associated with a number of cardiomyopathies in adult humans, they are in fact predominant in children [31]. Our data indicates that in adult zebrafish the majority of T waves are negative (albeit well below the threshold of what is considered abnormal for an adult human) (Figure 1C–E and Table 2) which may reflect differences in cardiac anatomy between humans and adult zebrafish. Taken together our data indicate that the most reliable technique for assessing the QT interval accurately is the exposed heart configuration.

### Effects of osmotic shock on adult zebrafish ECG characteristics

In humans, cerebral edema resulting from brain injury is frequently treated with hyperosmotic therapy to relieve inter cranial hypertension [32]. However, recent evidence suggests that elevating plasma osmolality can also lead to increased QTc and a higher risk of cardiac arrhythmias [33,34]. We therefore sought to determine what effect osmotic perturbations had on zebrafish ECG patterns. To achieve this we recorded ECGs in each of the 3 configurations in either hyperosmotic, isosmotic or hyposmotic conditions. Interestingly, osmotic challenge did not significantly alter the ECG characteristics in any of the three configurations when compared to the isosmotic controls (Table 3). However, despite the lack of significant differences in the ECG characteristics we were able to determine a clear positive QT-RR relationship in the exposed heart configuration which was not present in the unexposed configuration (Figure 2A–B). This positive correlation was also observed in the extracted heart configuration, however there was also a much higher variation of QT interval in relation to increasing RR in this configuration when compared to the exposed heart recordings (Figure 2B–C). Taken together our data indicate that although adult zebrafish can be a useful model for testing drug cardiotoxicity, due diligence should be taken when assessing therapies which can affect plasma osmolality as these may not affect zebrafish in the same way as humans. However it also appears that, unlike rodents, zebrafish hearts (like humans) elicit a positive QT-RR relationship allowing the QTc interval to be calculated which is vital when screening novel pharmaceuticals.

**Figure 2:**
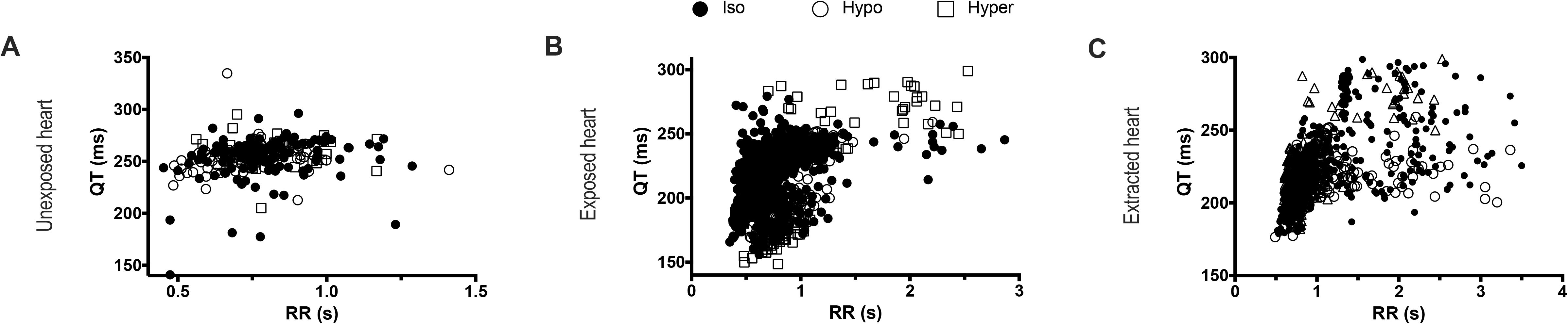
QT-RR relationship in different configurations of ECG measurement. **A**, unexposed heart (n=3); **B**, exposed heart (n=10); **C**, extracted heart (n=6). The QT-RR pairs, obtained in isosmotic, hypoosmotic and hyperosmotic conditions, are pooled.

### Effects of hypothermia on adult zebrafish ECG characteristics

Therapeutic hypothermia (a.k.a targeted temperature management) is often employed as neuroprotection in patients who have suffered a cardiac arrest or form other ischemic episodes resulting in reduced blood flow in the brain [35,36]. In order to ascertain whether zebrafish hearts respond to hypothermic conditions in a similar manner to humans we recorded ECGs in each configuration using either a control-warm (28°C) tyrode solution or a chilled (5°C) solution. In the unexposed heart configuration there was a significant reduction in the heart rate accompanied by significant elongations in the PR interval and QRS complex (Table 4). Although hypothermic conditions resulted in a lower heart rate in the exposed heart configuration this was below the threshold of significance. However, although the PR interval increased the QRS complex remained unchanged (Table 4). Importantly, as is the case in humans, hypothermia resulted in a significant lengthening of the QT interval in the exposed heart configuration (Table 4), which was also the case for the extracted heart configuration. These effects were all reversible (Table 4). Interestingly, hypothermia appears to increase the QT interval independently of the RR interval in both the exposed and extracted heart configurations (Figure 3B–C) indicating that this treatment has a direct effect on the QT interval. Taken together our data indicates that the exposed heart configuration appears to be the most reliable technique when using zebrafish to test hypothermic therapeutic treatments.

**Figure 3:**
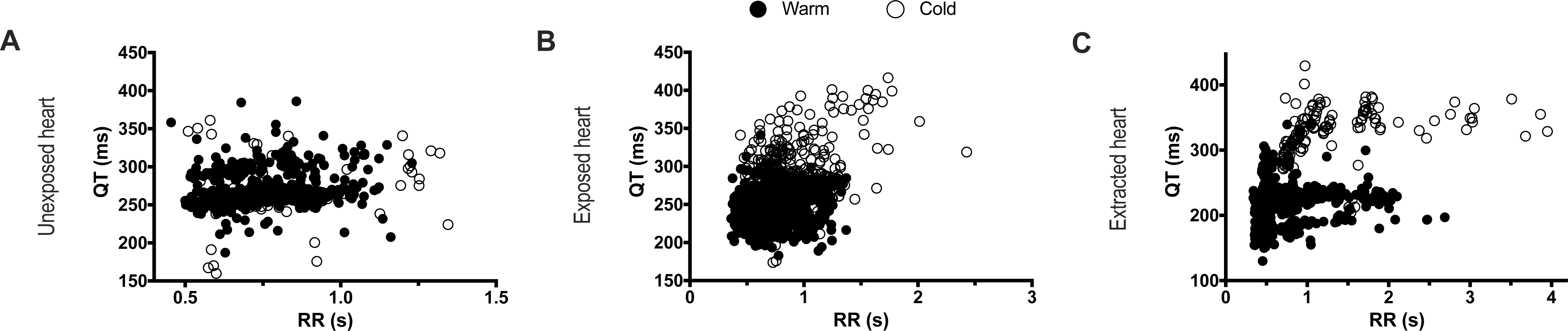
QT-RR relationship in different configurations of ECG measurement. **A**, unexposed heart (n=5); **B**, exposed heart (n=9); **C**, extracted heart (n=7). The QT-RR pairs, obtained in warm and cool conditions, are pooled.

### QT-RR relationship

Calculating the QTc is paramount during any drug cardiotoxicity testing. To achieve this it is necessary to establish the QT-RR relationship and in this sense identifying a clear T wave is essential. In the exposed heart configuration it is relatively easy to identify the T wave and thus build the QT-RR relationship. To characterize this relationship, we decided to use the Holzgrefe formula since it can be applied to different animal species if the Log(QT)-Log(RR) relationship is linear (QTch) [30]. When this is applied to the data obtained from the osmotic shock analysis we could observe a linear relationship with a slope of 0.2064 (Figure 4A). Based on this data we were subseqeuntly able to establish the non-linear fitting of the QT-RR relationship (Figure 4B). Using Holzgrefe’s formula we calculated the QTch (corrected holzgefe) and subsequently plotted the QTch-RR relationship (Figure 4C). It is apparent that there is no correlation between the QTch and the RR interval showing that QTch is independent of the RR interval as expected for a good correction (Figure 4C). Conversely, if we apply Bazett’s correction formula (QTcb) to our data, this transforms the positive QT-RR relationship into a negative QTcb-RR relationship (Figure 4C). Using this calculation, the QTcb is exaggeratedly increased by tachycardia (increased heart rate) and decreased by bradycardia (decreased heart rate), indicating that Holzgrefe’s formula provides the most accurate QTc value. To confirm that this was not due to our data, we re-analyzed previously published adult zebrafish ECG recordings [1] and fitted them with either the Holzgrefe or Bazett formula (Figure 4D). In this manner we were able to determine that applying the Holzgrefe formula results in a QTch that is independent of the RR interval, as observed with our own dataset. Conversely, applying the Bazett formula results in biased data similar to that which we observed with our own ECG recordings (Figure 4D). Taken together our data indicates that the exposed heart configuration is the most reliable for identifying T waves and establishing the QT-RR relationship. Furthermore, it is also apparent that Holzgrefe’s formula appears to be the most accurate method for calculating the corrected QT interval.

**Figure 4:**
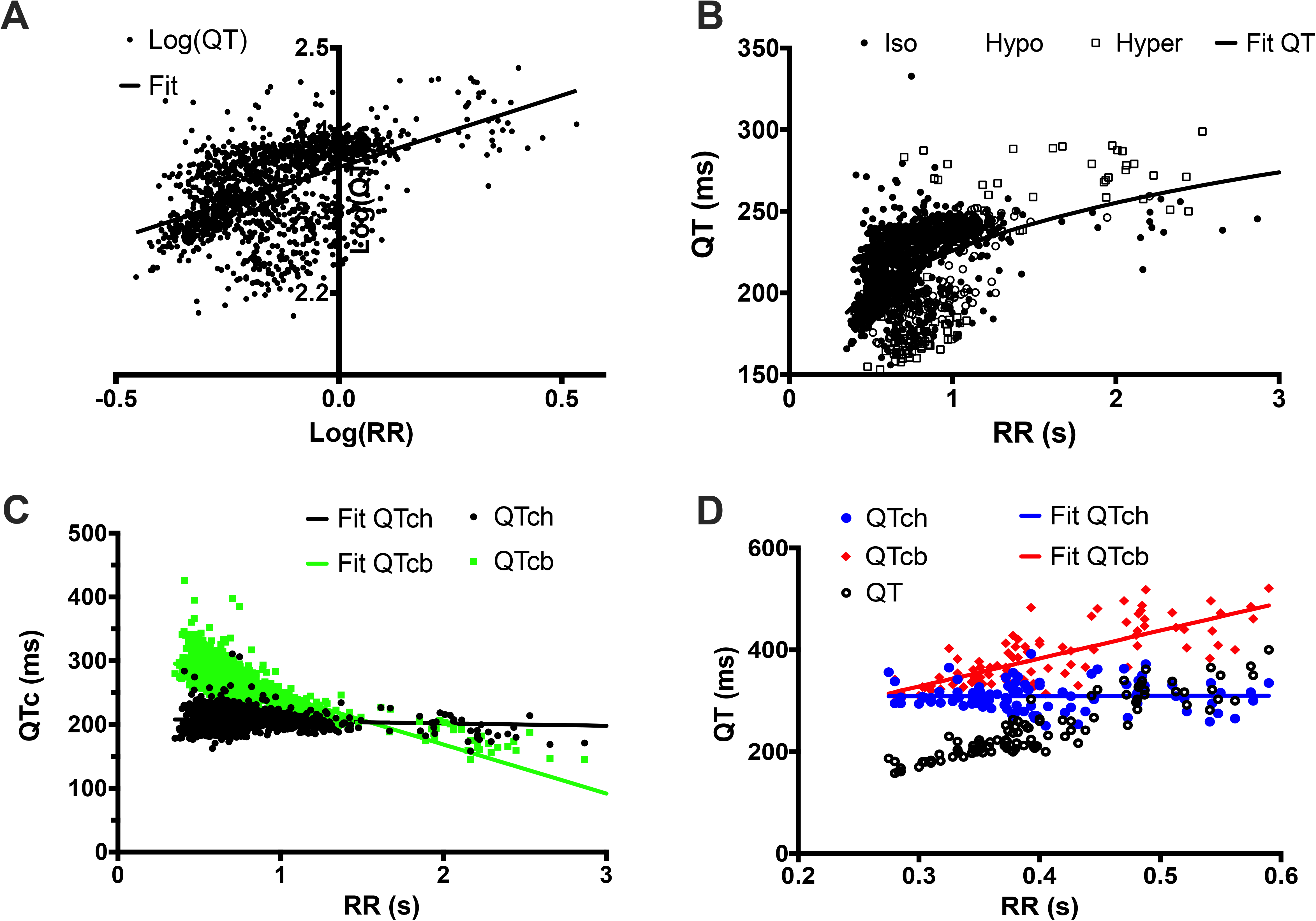
QT-RR relationship fitting. **A**, linear fitting (red line) of the Log(QT)-Log(RR) relationship (black dots). The slope of the relationship is 0.2064. **B**, non-linear fitting (red line) of the QT-RR relationship (black dots) using the characteristics of the fitting shown in A. **C**, QTc-RR relationship using Holzgrefe’s correction formula (QTch, black dots) and Bazett’s equation (QTcb, green dots). These relationships were linearly fitted (black line: QTch; green line: QTcb). **D**, QT-RR relationship from the data published in Milan *et al*., 2006. The QT-RR raw data are in open circles in black. The QT corrected by the Holzgrefe’s equation, QTch, are blue circles, and the corresponding linear fitting is the blue line while the QT corrected by Bazett’s formula are red diamonds and the corresponding linear fitting if the red line.

## Discussion

In this study, we have compared the ECG characteristics in three different configurations in order to evaluate their limitations and advantages. We have also assessed the effects of osmotic shock and hypothermia, two frequently used clinical treatments which are known to affect cardiac electrophysiology. By analyzing these data we have been able to better delineate the QT-RR relationship which is critical in calculating the QTc.

It is apparent from our own (and others [9,10]) research that recording adult zebrafish T waves is inherently difficult and susceptible to operator bias. Indeed, in the unexposed heart configuration we could barely detect any T waves at all. This is in contrast to the exposed heart configuration where we could readily observe distinct T waves and thus calculate the QT interval. Interestingly, adult zebrafish T waves can be either positive, as in healthy humans, or negative, as is in young children or other species such as canines. This phenomenon has also been previously described by Tsai *et al.* [24] who could detect a mixture of positive T waves (45%) and negative T waves (25%) (30% undetectable). From our own analysis in the exposed heart configuration (as opposed to the extracted heart configuration used by Tsai *et al.*) we observed similar levels of undetectable T waves (25%), however we found positive T waves in 20% of the recordings and negative T waves in 55% of the recordings. Furthermore, we observed that while the polarity of the T wave can be different between individual zebrafish it can also change within the same zebrafish during the experiment. This might be explained by the changing propagation of electrical gradient [12].

In humans hyperosmotic therapy is frequently used during the treatment of traumatic brain injuries. Recent evidence suggests that this treatment can also adversely affect the QT interval which could lead to potentially fatal cardiac arrhythmias. Our data indicates that, unlike humans, zebrafish cardiac electrophysiology is highly tolerant to changes in osmolality. Because zebrafish live in freshwater fish they are constantly subjected to the osmotic gradient between their environment and interstitial fluids. To prevent osmotic damage, zebrafish are able to rapidly regulate Na^+^ and Cl^−^ transport [37]. For example, a recent study from Kennard *et al*, showed that in zebrafish the process of wound healing was not affected by changes in osmolality despite considerable cell swelling [38]. The ability to adapt to osmotic shock, even after disrupting the epidermal barrier, was confirmed by our own results. This feature should also be taken into account when using zebrafish to perform cardiotoxicity tests of novel pharmaceuticals which may affect plasma osmolality or which may be used in combination with hyperosmotic therapy.

Following serious health complications, such as a heart attack or stroke, which result in a drastically reduced blood supply to the brain, target temperature management is often employed to reduce the risk of neuronal damage [39]. However this treatment is also known to impact cardiac electrophysiology and can lead to the lengthening of the QT interval and the potential for lethal arrhythmias [35,39]. Similarly, using the exposed heart configuration, we also observed that subjecting zebrafish to hypothermia resulted in a significant increase in the QT interval. This indicates that zebrafish would be an excellent model for testing the cardiotoxicity of novel therapies which cause or are used in conjunction with hypothermia.

Current legislation stipulates that before a new drug can be approved it must be demonstrated that it does not significantly alter the QT interval in human subjects. Clinical trials are very costly and before they can be undertaken pre-clinical assessment using animal models must be performed. The adult zebrafish is an excellent model for cardiotoxicity tests however it is imperative to first establish the QT-RR relationship accurately in order to ascertain whether any particular treatment actually affects the QT interval. The QT interval varies depending on the RR interval [1,2]. Thus it is important to correct the QT and make it independent of the heart rate when studying experimental conditions that change the RR. The standard equation used to correct the QT (QTc) is 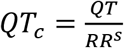. In humans, *s* equals 0.5 (Bazett’s equation) or 0.33 (Fridericia’s equation). These slope values have been widely used in correction formulae when studying zebrafish ECG recording [4,5,12–14,23,24,40]. However, it is vitally important to consider that the adult zebrafish mean cardiac frequency is around 120 bpm while in humans it is around 60 bpm. In this respect, we divided the RR interval by a zebrafish reference RR interval of 0.5s [30]. This correction allowed us to accurately calculate the QTc independently of the RR interval, something we could not achieve when employing Bazett’s formula.

Finally, the analysis of the QT-RR relationship raises some interesting questions. Indeed, compared to previous reports, we found a rather low *s* power of 0.18 (previously published s powers include-1.05 [1], 1.08 [3], 0.449 [41] and 0.58 [2]). This discrepancy between the QT-RR relationship maybe due to a number of factors such as the strain of fish [42], the experimental conditions, anesthesia and other types of treatment. Recently, other confounding factors such as sex and body weight have also been described [43]. This suggests that the best protocol when studying zebrafish ECG is to first characterize the QT-RR of a given strain of fish in standardized experimental conditions before going further. We would also like to highlight the risk of bias induced by correction formula such as Bazett’s equation which appears to be critical. Our data suggest that Holzgrefe’s formula provides the most accurate correction since the QTc does not change in relation to the RR. In contrast Bazett’s equation produces a negative relationship. This is important because under conditions that can reduce the cardiac rate it would appear that QTc interval would increase, which in fact would just be an artefact of the correction formula. In order to confirm this finding we utilized previously published QT-RR data [1] and we applied either Holzgrefe’s correction formula or the Bazett formula. In this manner we found that Bazett’s formula does not correct the QT, it even overestimates it at higher heart rates, while Holzgrefe’s formula perfectly corrected the QT. Thus, it is clear that Bazett’s formula has to be avoided to correct the QT interval in zebrafish. Furthermore, because Holzgrefe’s correction is made relative to the animals heart rate this allows the QTc to be calculated in the range of the measured QT which is not the case without this normalization.

In conclusion, of the three commonly used techniques it seems that the exposed heart configuration provides the most reliable ECG recordings in adult zebrafish. Furthermore, despite the considerable evolutionary distance between zebrafish and humans their ECG characteristics are remarkably similar which highlights the utility of using adult zebrafish for cardiotoxicity tests over other animal models such as rodents.

## Acknowledgments

We thank Arnaud Gac (AD Instrument) for helping in the setting and improvement of the ECG apparatus system. The Jopling lab (EA, LR, JB and CJ) is part of the Laboratory of Excellence Ion Channel Science and Therapeutics supported by a grant from the ANR. Work in the Jopling lab is also supported by a grant from the “la Fondation Leducq”.

## Authors contribution

EA, LR and JYL performed the experiments. EA, LR, JYL, EB, AGT, JB, JT, CJ and MD participated to the design of the experiments, discussed the results and participated in writing the manuscript.

## References

1. Milan, D.J.; Jones, I.L.; Ellinor, P.T.; MacRae, C.A. In Vivo Recording of Adult Zebrafish Electrocardiogram and Assessment of Drug-Induced QT Prolongation. Am J Physiol Heart Circ Physiol 2006, 291, H269–273, doi:10.1152/ajpheart.00960.2005.

2. Arnaout, R.; Ferrer, T.; Huisken, J.; Spitzer, K.; Stainier, D.Y.R.; Tristani-Firouzi, M.; Chi, N.C. Zebrafish Model for Human Long QT Syndrome. Proc Natl Acad Sci U S A 2007, 104, 11316–11321, doi:10.1073/pnas.0702724104.

3. Chaudhari, G.H.; Chennubhotla, K.S.; Chatti, K.; Kulkarni, P. Optimization of the Adult Zebrafish ECG Method for Assessment of Drug-Induced QTc Prolongation. J Pharmacol Toxicol Methods 2013, 67, 115–120, doi:10.1016/j.vascn.2013.01.007.

4. Liu, C.C.; Li, L.; Lam, Y.W.; Siu, C.W.; Cheng, S.H. Improvement of Surface ECG Recording in Adult Zebrafish Reveals That the Value of This Model Exceeds Our Expectation. Sci Rep 2016, 6, 25073, doi:10.1038/srep25073.

5. Lin, M.-H.; Chou, H.-C.; Chen, Y.-F.; Liu, W.; Lee, C.-C.; Liu, L.Y.-M.; Chuang, Y.-J. Development of a Rapid and Economic in Vivo Electrocardiogram Platform for Cardiovascular Drug Assay and Electrophysiology Research in Adult Zebrafish. Sci Rep 2018, 8, 15986, doi:10.1038/s41598-018-33577-7.

6. Echeazarra, L.; Hortigón-Vinagre, M.P.; Casis, O.; Gallego, M. Adult and Developing Zebrafish as Suitable Models for Cardiac Electrophysiology and Pathology in Research and Industry. Front Physiol 2020, 11, 607860, doi:10.3389/fphys.2020.607860.

7. Nemtsas, P.; Wettwer, E.; Christ, T.; Weidinger, G.; Ravens, U. Adult Zebrafish Heart as a Model for Human Heart? An Electrophysiological Study. J Mol Cell Cardiol 2010, 48, 161–171, doi:10.1016/j.yjmcc.2009.08.034.

8. Brette, F.; Luxan, G.; Cros, C.; Dixey, H.; Wilson, C.; Shiels, H.A. Characterization of Isolated Ventricular Myocytes from Adult Zebrafish (Danio Rerio). Biochem Biophys Res Commun 2008, 374, 143–146, doi:10.1016/j.bbrc.2008.06.109.

9. Brönnimann, D.; Djukic, T.; Triet, R.; Dellenbach, C.; Saveljic, I.; Rieger, M.; Rohr, S.; Filipovic, N.; Djonov, V. Pharmacological Modulation of Hemodynamics in Adult Zebrafish In Vivo. PLoS One 2016, 11, e0150948, doi:10.1371/journal.pone.0150948.

10. Le, T.; Lenning, M.; Clark, I.; Bhimani, I.; Fortunato, J.; Mash, P.; Xu, X.; Cao, H. Acquisition, Processing and Analysis of Electrocardiogram in Awake Zebrafish. IEEE Sens J 2019, 19, 4283–4289, doi:10.1109/jsen.2019.2897789.

11. Song, J.; Qiao, L.; Ji, L.; Ren, B.; Hu, Y.; Zhao, R.; Ren, Z. Toxic Responses of Zebrafish (Danio Rerio) to Thallium and Deltamethrin Characterized in the Electrocardiogram. Chemosphere 2018, 212, 1085–1094, doi:10.1016/j.chemosphere.2018.09.014.

12. Zhao, Y.; James, N.A.; Beshay, A.R.; Chang, E.E.; Lin, A.; Bashar, F.; Wassily, A.; Nguyen, B.; Nguyen, T.P. Adult Zebrafish Ventricular Electrical Gradients as Tissue Mechanisms of ECG Patterns under Baseline vs. Oxidative Stress. Cardiovasc Res 2021, 117, 1891–1907, doi:10.1093/cvr/cvaa238.

13. Yu, F.; Li, R.; Parks, E.; Takabe, W.; Hsiai, T.K. Electrocardiogram Signals to Assess Zebrafish Heart Regeneration: Implication of Long QT Intervals. Ann Biomed Eng 2010, 38, 2346–2357, doi:10.1007/s10439-010-9993-6.

14. Chablais, F.; Veit, J.; Rainer, G.; Jaźwińska, A. The Zebrafish Heart Regenerates after Cryoinjury-Induced Myocardial Infarction. BMC Dev Biol 2011, 11, 21, doi:10.1186/1471-213X-11-21.

15. Lee, J.; Cao, H.; Kang, B.J.; Jen, N.; Yu, F.; Lee, C.-A.; Fei, P.; Park, J.; Bohlool, S.; Lash-Rosenberg, L.; et al. Hemodynamics and Ventricular Function in a Zebrafish Model of Injury and Repair. Zebrafish 2014, 11, 447–454, doi:10.1089/zeb.2014.1016.

16. Zhang, X.; Tai, J.; Park, J.; Tai, Y.-C. Flexible MEA for Adult Zebrafish ECG Recording Covering Both Ventricle and Atrium. In Proceedings of the 2014 IEEE 27th International Conference on Micro Electro Mechanical Systems (MEMS); January 2014; pp. 841–844.

17. Lenning, M.; Fortunato, J.; Le, T.; Clark, I.; Sherpa, A.; Yi, S.; Hofsteen, P.; Thamilarasu, G.; Yang, J.; Xu, X.; et al. Real-Time Monitoring and Analysis of Zebrafish Electrocardiogram with Anomaly Detection. Sensors (Basel) 2017, 18, E61, doi:10.3390/s18010061.

18. Zhao, Y.; Yun, M.; Nguyen, S.A.; Tran, M.; Nguyen, T.P. In Vivo Surface Electrocardiography for Adult Zebrafish. J Vis Exp 2019, doi:10.3791/60011.

19. Sun, P.; Zhang, Y.; Yu, F.; Parks, E.; Lyman, A.; Wu, Q.; Ai, L.; Hu, C.-H.; Zhou, Q.; Shung, K.; et al. Micro-Electrocardiograms to Study Post-Ventricular Amputation of Zebrafish Heart. Ann Biomed Eng 2009, 37, 890–901, doi:10.1007/s10439-009-9668-3.

20. Zhang, X.; Beebe, T.; Jen, N.; Lee, C.-A.; Tai, Y.; Hsiai, T.K. Flexible and Waterproof Micro-Sensors to Uncover Zebrafish Circadian Rhythms: The next Generation of Cardiac Monitoring for Drug Screening. Biosens Bioelectron 2015, 71, 150–157, doi:10.1016/j.bios.2015.04.027.

21. Yu, F.; Zhao, Y.; Gu, J.; Quigley, K.L.; Chi, N.C.; Tai, Y.-C.; Hsiai, T.K. Flexible Microelectrode Arrays to Interface Epicardial Electrical Signals with Intracardial Calcium Transients in Zebrafish Hearts. Biomed Microdevices 2012, 14, 357–366, doi:10.1007/s10544-011-9612-9.

22. Cao, H.; Yu, F.; Zhao, Y.; Zhang, X.; Tai, J.; Lee, J.; Darehzereshki, A.; Bersohn, M.; Lien, C.-L.; Chi, N.C.; et al. Wearable Multi-Channel Microelectrode Membranes for Elucidating Electrophysiological Phenotypes of Injured Myocardium. Integr Biol (Camb) 2014, 6, 789–795, doi:10.1039/c4ib00052h.

23. Meder, B.; Scholz, E.P.; Hassel, D.; Wolff, C.; Just, S.; Berger, I.M.; Patzel, E.; Karle, C.; Katus, H.A.; Rottbauer, W. Reconstitution of Defective Protein Trafficking Rescues Long-QT Syndrome in Zebrafish. Biochem Biophys Res Commun 2011, 408, 218–224, doi:10.1016/j.bbrc.2011.03.121.

24. Tsai, C.-T.; Wu, C.-K.; Chiang, F.-T.; Tseng, C.-D.; Lee, J.-K.; Yu, C.-C.; Wang, Y.-C.; Lai, L.-P.; Lin, J.-L.; Hwang, J.-J. In-Vitro Recording of Adult Zebrafish Heart Electrocardiogram - a Platform for Pharmacological Testing. Clin Chim Acta 2011, 412, 1963–1967, doi:10.1016/j.cca.2011.07.002.

25. Stoyek, M.R.; Quinn, T.A.; Croll, R.P.; Smith, F.M. Zebrafish Heart as a Model to Study the Integrative Autonomic Control of Pacemaker Function. Am J Physiol Heart Circ Physiol 2016, 311, H676–688, doi:10.1152/ajpheart.00330.2016.

26. Schwerte, T.; Prem, C.; Mairösl, A.; Pelster, B. Development of the Sympatho-Vagal Balance in the Cardiovascular System in Zebrafish (Danio Rerio) Characterized by Power Spectrum and Classical Signal Analysis. J Exp Biol 2006, 209, 1093–1100, doi:10.1242/jeb.02117.

27. Stoyek, M.R.; Hortells, L.; Quinn, T.A. From Mice to Mainframes: Experimental Models for Investigation of the Intracardiac Nervous System. J Cardiovasc Dev Dis 2021, 8, 149, doi:10.3390/jcdd8110149.

28. Champeroux, P.; Le Guennec, J.Y.; Jude, S.; Laigot, C.; Maurin, A.; Sola, M.L.; Fowler, J.S.L.; Richard, S.; Thireau, J. The High Frequency Relationship: Implications for Torsadogenic HERG Blockers. Br J Pharmacol 2016, 173, 601–612, doi:10.1111/bph.13391.

29. Champéroux, P.; Fesler, P.; Judé, S.; Richard, S.; Le Guennec, J.-Y.; Thireau, J. High-Frequency Autonomic Modulation: A New Model for Analysis of Autonomic Cardiac Control. Br J Pharmacol 2018, 175, 3131–3143, doi:10.1111/bph.14354.

30. Holzgrefe, H.; Ferber, G.; Champeroux, P.; Gill, M.; Honda, M.; Greiter-Wilke, A.; Baird, T.; Meyer, O.; Saulnier, M. Preclinical QT Safety Assessment: Cross-Species Comparisons and Human Translation from an Industry Consortium. J Pharmacol Toxicol Methods 2014, 69, 61–101, doi:10.1016/j.vascn.2013.05.004.

31. Dickinson, D.F. The Normal ECG in Childhood and Adolescence. Heart 2005, 91, 1626–1630, doi:10.1136/hrt.2004.057307.

32. Raslan, A.; Bhardwaj, A. Medical Management of Cerebral Edema. Neurosurg Focus 2007, 22, E12, doi:10.3171/foc.2007.22.5.13.

33. Dabrowski, W.; Siwicka-Gieroba, D.; Robba, C.; Badenes, R.; Bialy, M.; Iwaniuk, P.; Schlegel, T.T.; Jaroszynski, A. Plasma Hyperosmolality Prolongs QTc Interval and Increases Risk for Atrial Fibrillation in Traumatic Brain Injury Patients. J Clin Med 2020, 9, E1293, doi:10.3390/jcm9051293.

34. Dabrowski, W.; Siwicka-Gieroba, D.; Robba, C.; Bielacz, M.; Sołek-Pastuszka, J.; Kotfis, K.; Bohatyrewicz, R.; Jaroszyński, A.; Malbrain, M.L.N.G.; Badenes, R. Potentially Detrimental Effects of Hyperosmolality in Patients Treated for Traumatic Brain Injury. J Clin Med 2021, 10, 4141, doi:10.3390/jcm10184141.

35. Khan, J.N.; Prasad, N.; Glancy, J.M. QTc Prolongation during Therapeutic Hypothermia: Are We Giving It the Attention It Deserves? Europace 2010, 12, 266–270, doi:10.1093/europace/eup376.

36. Rosol, Z.; Miranda, D.F.; Sandoval, Y.; Bart, B.A.; Smith, S.W.; Goldsmith, S.R. The Effect of Targeted Temperature Management on QT and Corrected QT Intervals in Patients with Cardiac Arrest. J Crit Care 2017, 39, 182–184, doi:10.1016/j.jcrc.2017.02.030.

37. Boisen, A.M.Z.; Amstrup, J.; Novak, I.; Grosell, M. Sodium and Chloride Transport in Soft Water and Hard Water Acclimated Zebrafish (Danio Rerio). Biochim Biophys Acta 2003, 1618, 207–218, doi:10.1016/j.bbamem.2003.08.016.

38. Kennard, A.S.; Theriot, J.A. Osmolarity-Independent Electrical Cues Guide Rapid Response to Injury in Zebrafish Epidermis. Elife 2020, 9, e62386, doi:10.7554/eLife.62386.

39. Jandu, S.; Sefa, N.; Sawyer, K.N.; Swor, R. Electrocardiographic Changes in Patients Undergoing Targeted Temperature Management. J Am Coll Emerg Physicians Open 2020, 1, 327–332, doi:10.1002/emp2.12104.

40. Hassel, D.; Scholz, E.P.; Trano, N.; Friedrich, O.; Just, S.; Meder, B.; Weiss, D.L.; Zitron, E.; Marquart, S.; Vogel, B.; et al. Deficient Zebrafish Ether-à-Go-Go-Related Gene Channel Gating Causes Short-QT Syndrome in Zebrafish Reggae Mutants. Circulation 2008, 117, 866–875, doi:10.1161/CIRCULATIONAHA.107.752220.

41. Mersereau, E.J.; Poitra, S.L.; Espinoza, A.; Crossley, D.A.; Darland, T. The Effects of Cocaine on Heart Rate and Electrocardiogram in Zebrafish (Danio Rerio). Comp Biochem Physiol C Toxicol Pharmacol 2015, 172–173, 1–6, doi:10.1016/j.cbpc.2015.03.007.

42. Holden, L.A.; Brown, K.H. Baseline MRNA Expression Differs Widely between Common Laboratory Strains of Zebrafish. Sci Rep 2018, 8, 4780, doi:10.1038/s41598-018-23129-4.

43. Duong, T.; Rose, R.; Blazeski, A.; Fine, N.; Woods, C.E.; Thole, J.F.; Sotoodehnia, N.; Soliman, E.Z.; Tung, L.; McCallion, A.S.; et al. Development and Optimization of an in Vivo Electrocardiogram Recording Method and Analysis Program for Adult Zebrafish. Dis Model Mech 2021, 14, dmm048827, doi:10.1242/dmm.048827.

