## Supplementary material for "Characterization of the adult zebrafish electrocardiogram": Tables

**Table 1**

| Configurations | PR (ms) | QRS (ms) | QT (ms) | RR (s) | HR (bpm) |
|---|---|---|---|---|---|
| **Unexposed heart** | 44.79 (1.90) *(20)* | 25.93 (1.02) *(20)* | 273.4 (7.65) *(8)* | 766.8 (30.92) *(20)* | 80.27 (3.07) *(20)* |
| **Exposed heart** | 41.44 (2.22) *(17)* | 31.34 (1.54)* *(20)* | 234.8 (6.29) *(16)* | 699.9 (32.99) *(20)* | 86.46 (4.40) *(20)* |
| **Extracted heart** | 49.87 (6.52) *(5)* | 34.09 (4.94) *(17)* | 213.4 (17.60)$$ *(13)* | 910.1 (85.80) *(18)* | 74.68 (6.24) *(18)* |

Table 1: Characteristics of the ECG in the different conditions of recording. Data are presented as mean (SEM) *(number of fish)*. One-way ANOVA (or Kruskal-Wallis test), followed by an appropriate *post hoc* test were performed to compare the effect of recording conditions on the different ECG parameters. Statistical differences (p<0.05) are indicated by a * when values are different from the Unexposed heart configuration, $ when values are different from the Exposed heart. The QT intervals of Unexposed heart were not statistically analyzed. NC: not calculable.

**Table 2**

| Configurations | Positive T wave | Negative T wave | Undetectable |
|---|---|---|---|
| **Unexposed Heart** | 0% (0/20) | 40% (8/20) | 60% (12/20) |
| **Exposed Heart** | 25% (5/20) | 55% (11/20) | 20% (4/20) |
| **Extracted Heart** | 28% (5/18) | 44% (8/18) | 28% (5/18) |

Table 2: Characteristics of the T waves in the different recording conditions. Data are presented as percentages (proportion of animals).

**Table 3**

| Configuration / Condition | PR (ms) | QRS (ms) | QT (ms) | RR (s) | HR (bpm) |
|---|---|---|---|---|---|
| **Unexposed heart** | | | | | |
| **Tyrode** | 46.15 (3.34) *(10)* | 26.65 (1.24) *(10)* | 271.5 (9.01) *(3)* | 799.5 (41.08) *(10)* | 76.82 (4.01) *(10)* |
| **Isosmotic** | 57.01 (13.14) *(10)* | 25.86 (1.45) *(10)* | 265.1 (8.38) *(3)* | 756.5 (44.95) *(10)* | 81.34 (3.89) *(10)* |
| **Hyposmotic** | 47.08 (2.33) *(9)* | 26.11 (1.47) *(10)* | 266.8 (4.24) *(3)* | 727.7 (46.7) *(10)* | 84.85 (4.25) *(10)* |
| **Wash isosmotic** | 46.97 (1.95) *(7)* | 24.86 (1.79) *(8)* | NC *(1)* | 743.3 (38.04) *(8)* | 82.27 (4.36) *(8)* |
| **Hyperosmotic** | 46.59 (2.44) *(8)* | 25.95 (1.22) *(10)* | NC *(2)* | 766.2 (44.61) *(10)* | 80.50 (4.26) *(10)* |
| **Exposed heart** | | | | | |
| **Tyrode** | 43.78 (4.04) *(10)* | 33.80 (1.46) *(10)* | 235.0 (9.99) *(10)* | 705.4 (45.32) *(10)* | 88.48 (5.86) *(10)* |
| **Isosmotic** | 42.13 (2.02) *(9)* | 31.39 (1.47) *(10)* | 241.3 (11.73) *(8)* | 733.2 (149.0) *(10)* | 85.3 (6.08) *(10)* |
| **Hyposmotic** | 41.07 (2.02) *(9)* | 29.27 (1.69) *(10)* | 244.9 (12.9) *(7)* | 716.6 (46.9) *(10)* | 87.43 (6.37) *(10)* |
| **Wash isosmotic** | 41.36 (1.98) *(9)* | 29.39 (1.74) *(10)* | 243.9 (14.02) *(6)* | 710.4 (51.48) *(10)* | 88.86 (6.92) *(10)* |
| **Hyperosmotic** | 40.70 (1.83) *(9)* | 31.07 (1.55) *(10)* | 245.2 (14.48) *(5)* | 729.6 (48.69) *(10)* | 85.88 (6.18) *(10)* |
| **Extracted heart** | | | | | |
| **Tyrode** | NC *(0)* | 42.82 (8.06) *(9)* | 233.9 (19.11) *(6)* | 954.0 (102.5) *(10)* | 69.79 (7.80) *(10)* |
| **Isosmotic** | NC *(0)* | 33.23 (5.01) *(9)* | 232.2 (6.18) *(5)* | 1239.0 (240.0) *(10)* | 67.24 (12.14) *(10)* |
| **Hyposmotic** | NC *(0)* | 32.00 (4.84) *(9)* | 224.4 (6.71) *(7)* | 1201.0 (202.6) *(10)* | 63.27 (9.95) *(10)* |
| **Wash isosmotic** | NC *(0)* | 31.99 (5.05) *(9)* | 215.7 (6.17) *(7)* | 980.7 (180.7) *(10)* | 77.25 (10.31) *(10)* |
| **Hyperosmotic** | NC *(0)* | 31.18 (4.14) *(8)* | 227.7 (8.44) *(7)* | 1132.0 (186.0) *(10)* | 64.46 (8.23) *(10)* |

Table 3: Effects of osmotic shocks on the characteristics of the ECG. Data are presented as mean (SEM) *(number of fish)*. One-way ANOVA (or Kruskal-Wallis test), followed by an appropriate *post hoc* test were performed to compare osmotic conditions within each group. Statistical differences ($p<0.05$) are indicated by a * when values are different from the Tyrode condition, $^{\$}$ when values are different from the isosmotic. The QT intervals of Unexposed heart were not statistically analyzed. NC: not calculable.

**Table 4**

| Configuration Condition | PR (ms) | QRS (ms) | QT (ms) | RR (s) | HR (bpm) |
|---|---|---|---|---|---|
| **Unexposed heart** | | | | | |
| **Warm (control)** | 43.42 (1.90) *(10)* | 25.22 (1.66) *(10)* | 273.4 (7.65) *(5)* | 734 (45.94) *(10)* | 83.72 (4.60) *(10)* |
| **Cold** | 78.47 (6.87)**** *(10)* | 37.59 (2.71)** *(10)* | 252.2 (43.08) *(2)* | 1401 (196.9)** *(10)* | 50.86 (7.30)** *(10)* |
| **Warm** | 46.95 (2.23)$$$ *(8)* | 25.33 (2.11)$$ *(9)* | 272.5 (26.66) *(3)* | 921.4 (81.02)$ (10) | 69.05 (5.84) *(10)* |
| **Exposed heart** | | | | | |
| **Warm (control)** | 38.48 (1.91) *(8)* | 28.88 (2.55) *(10)* | 234.5 (4.07) *(6)* | 694.4 (50.34) *(10)* | 84.44 (6.80) *(10)* |
| **Cold** | 49.78 (2.96)** *(9)* | 29.96 (2.35) *(10)* | 292.6 (11.86)** *(9)* | 956.6 (76.04)* *(10)* | 65.36 (4.43) *(10)* |
| **Warm** | 41.32 (1.94)$ *(9)* | 25.99 (2.96 *(10)* | 244.5 (6.57)$$ *(8)* | 762.9 (74.31) *(10)* | 83.74 (76) *(10)* |
| **Extracted heart** | | | | | |
| **Warm (control)** | 49.87 (6.52) *(5)* | 24.26 (3) *(8)* | 195.8 (27.90) *(7)* | 855.1 (150.3) *(8)* | 80.79 (10.29) *(8)* |
| **Cold** | 56.23 (6.65) *(4)* | 38.80 (3.92) *(8)* | 341.8 (9.05)* *(5)* | 2038 (462.5)* *(8)* | 39.28 (6.97)* *(8)* |
| **Warm** | 41.42 (3.45) *(3)* | 34.52 (11.95) *(8)* | 220.5 (11.62)$ *(7)* | 846 (116.9)$ *(8)* | 77.68 (10.08)$ (8) |

Table 4: Effects of temperature on the characteristics of the ECG. Data are presented as mean (SEM) *(number of fish)*. One-way ANOVA (or Kruskal-Wallis test), followed by an appropriate *post hoc* test were performed to compare temperature conditions within each group. Statistical differences ($p<0.05$) are indicated by a * when values are different from the Warm (control) condition, $ when values are different from the cold condition. The QT intervals of Unexposed heart were not statistically analyzed. NC: not calculable.

**Supplemental Table 1**

| Wave detection / Condition | P wave (%) | T wave (%) |
|---|---|---|
| **Unexposed heart** | | |
| **Tyrode** | 94.1% (1.9) *(10)* | 45.6% (14.7) *(5)* |
| **Isosmotic** | 78.4% (8.7) *(9)* | 56.4% (18.0) *(5)* |
| **Hyposmotic** | 76.2% (10.5) *(9)* | 40.4% (16.2) *(5)* |
| **Wash isosmotic** | 79.0% (10.0) *(7)* | 26.7% (17.4) *(3)* |
| **Hyperosmotic** | 79.0% (8.8) *(8)* | 39.8% (17.9) *(4)* |
| **Exposed heart** | | |
| **Tyrode** | 91.2% (8.5) *(10)* | 90.8% (3.2) *(10)* |
| **Isosmotic** | 98.3% (0.7) *(9)* | 86.9% (4.4) *(8)* |
| **Hyposmotic** | 98.8% (0.5) *(9)* | 84.0% (7.2) *(6)* |
| **Wash isosmotic** | 98.2% (1.2) *(9)* | 82.2% (6.5) *(6)* |
| **Hyperosmotic** | 99.7% (0.2) *(9)* | 92.6% (2.5) *(5)* |
| **Extracted heart** | | |
| **Tyrode** | NC (NC) *(0)* | 60.1% (15.3) *(7)* |
| **Isosmotic** | NC (NC) *(0)* | 73.5% (10.5) *(8)* |
| **Hyposmotic** | NC (NC) *(0)* | 86.6% (4.0) *(7)* |
| **Wash isosmotic** | NC (NC) *(0)* | 77.1% (11.6) *(8)* |
| **Hyperosmotic** | NC (NC) *(0)* | 92.1% (5.0) *(7)* |

Supplemental Table 1: P and T waves detection in the osmolarity group. Proportion of ECG complexes presenting P/T wave during a 1-minute recording. Data are presented as mean (SEM) *(number of fish)*.

### Supplemental Table 2

| Wave detection<br>Condition | P wave (%) | T wave (%) |
|---|---|---|
| **Unexposed heart** | | |
| **Warm (control)** | 86.4% (6.7)<br>*(10)* | 62.6% (8.8)<br>*(5)* |
| **Cold** | 59.4% (10.5)<br>*(10)* | NC (NC)<br>*(1)* |
| **Warm** | 68.2% (10.6)<br>*(9)* | 17.3% (0.7)<br>*(3)* |
| **Exposed heart** | | |
| **Warm (control)** | 75.4% (32.8)<br>*(8)* | 96.5% (1.6)<br>*(6)* |
| **Cold** | 80.7% (7.9)<br>*(9)* | 71.9% (7.3)<br>*(9)* |
| **Warm** | 91.8% (3.1)<br>*(9)* | 82.5% (5.9)<br>*(8)* |
| **Extracted heart** | | |
| **Warm (control)** | 62.0% (14.9)<br>*(7)* | 61.3% (9.6)<br>*(8)* |
| **Cold** | 85.0% (10.2)<br>*(4)* | 68.2% (13.4)<br>*(6)* |
| **Warm** | 59.5% (22.5)<br>*(4)* | 58.4% (11.2)<br>*(7)* |

Supplemental Table 2: P and T waves detection in the temperature group. Proportion of ECG complexes presenting P/T wave during a 1-minute recording. Data are presented as mean (SEM) *(number of fish)*.
